# Body mass aging trajectory is modulated by environmental conditions but independent of lifespan

**DOI:** 10.1101/376327

**Authors:** Michael Briga, Blanca Jimeno, Simon Verhulst

**Author notes:** Department of Biology, University of Turku, Turku, Finland (Michael Briga).

## Abstract

How lifespan associates with aging trajectories of health and disease is an urgent question in societies with increasing lifespan. Body mass declines with age are associated with decreased organismal functioning in many species. We tested whether two factors that decreased lifespan in zebra finches, sex and manipulated environmental quality, accelerated the onset and/or rate of within-individual body mass declines. We subjected 597 birds for nine years to experimentally manipulated foraging costs (harsh = H, benign = B) during development and in adulthood in a 2×2 design. This yielded four treatment combinations (HH, HB, BH, BB). Harsh environments during development and in adulthood decreased average body mass additively. In males, the aging trajectory was quadratic, with a maximum between 3.5 and 4 years, and independent of the environment (HH=HB=BH=BB). In females, the shape of the aging trajectory differed between environments: a quadratic trajectory as in males in the benign adult environment (HB=BB), a linear decline when benign development was followed by harsh adulthood (BH) and a linear increase when in a lifelong harsh environment (HH). We found no evidence for an association between lifespan and body mass aging trajectories either between or within experimental groups. However, females lived shorter than males, and their body mass decline started earlier for most treatment combinations. Thus, we conclude that foraging conditions can affect the shape of body mass aging trajectories, but these are independent of lifespan.

## Introduction

Senescence is the decline in organismal functioning with age resulting in declining fecundity and survival. Aging is a change in trait functioning with age, which may or may not be associated with declines in fecundity or survival. Aging ends in death and therefore the (implicit) assumption is often made that factors that changes in lifespan also alter aging. However, aging can differ from lifespan in that it explicitly refers to the *change* in organismal functioning preceding death. Hence to what extent factors that alter lifespan also alter aging remains to be identified (Bansal et al., 2015; Christensen et al., 2009; Hansen and Kennedy, 2016; Williams, 1999). This issue is of major relevance to contemporary society. Life expectancy has increased continuously since the 19^th^ century, but to what extent this increase is accompanied by delays in aging remains unclear (Christensen et al., 2009). Hence, to what extent aging and lifespan are scaled and affected by the same factors remains an issue in our ever longer-living society.

A key point underlying the association between aging and lifespan is identifying how organisms and traits change with age. Aging can occur following a variety of trajectories, which can be characterized by an age of onset, a rate and a shape. It is known that environmental quality during development and/or adulthood can alter the onset and rate of aging (Bouwhuis et al., 2010; Lemaitre and Gaillard, 2017; Nussey et al., 2013, 2007). In contrast, what determines the shape of the aging trajectory remains poorly known. Aging shapes can vary widely (Fig. 1). For example, aging may start at a certain age and decline linearly (Fig. 1A) or accelerating till death (Fig. 1B). This accelerating scenario was described for body mass in humans (Kuk et al., 2009) and laboratory rodents (Miller et al., 2002; Murtagh-Mark et al., 1995; Yu et al., 1985). Alternatively, aging may occur sharply before death, a phenomenon coined terminal decline (Fig. 1C). This scenario was described for a variety of traits in birds, including social dominance, sexual signals, telomere length and reproduction (Coulson and Fairweather, 2001; Rattiste, 2004; Salomons et al., 2009; Simons et al., 2016; Torres et al., 2011; Verhulst et al., 2014). The shape of aging trajectories are often described as trait-specific (Gaillard and Lemaitre, 2017; Hayward et al., 2015). Unfortunately, individual variation in aging shapes has rarely been investigated. Hence, what determines aging shapes and to what extent they are individual-specific remains poorly known.

**Fig. 1:**
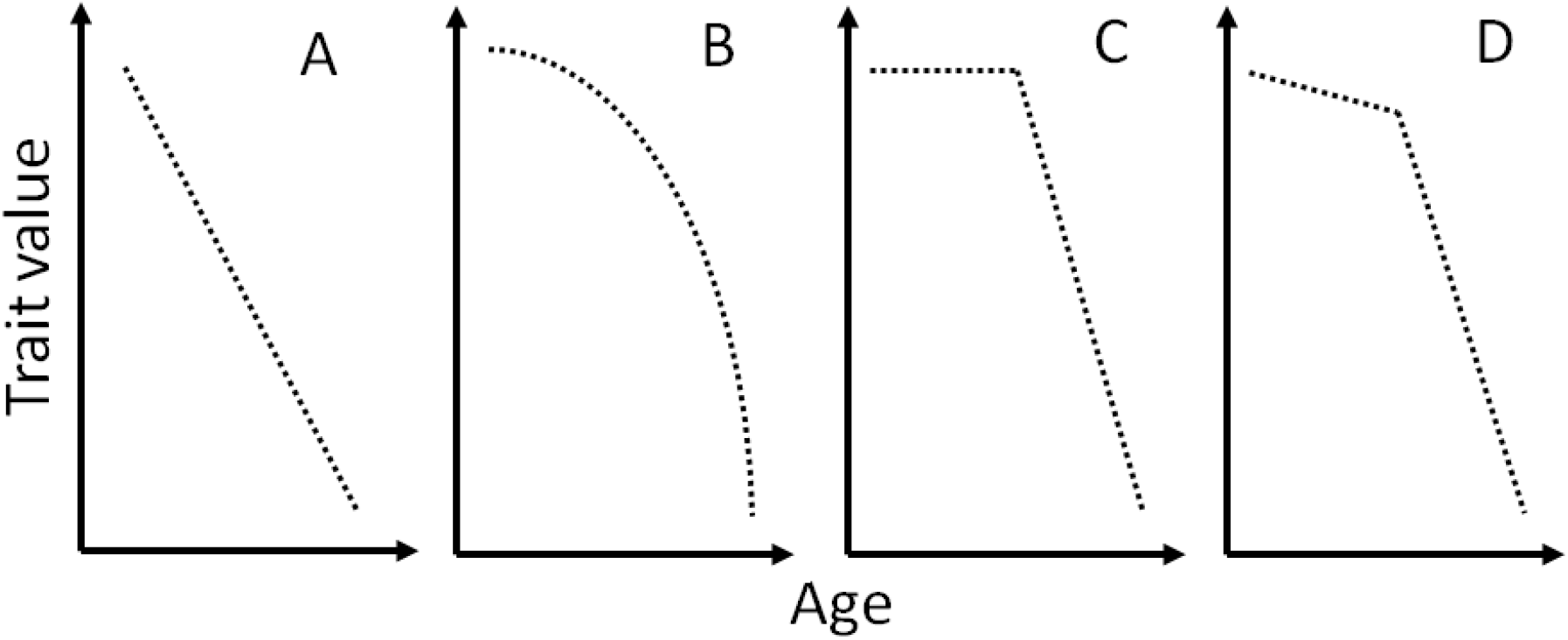
Schematic representation of four aging shapes tested in this manuscript, i.e. starting from the age at which traits decline in performance. Aging may be determined by chronological age, following gradual (A) or accelerating decline (B). Alternatively, aging may be better described by years before death resulting in a terminal decline (C) or by a combination of a chronological and terminal decline (D).

Here, we test whether factors that affect lifespan concomitantly alter body mass aging in zebra finches. Body mass predicts survival or lifespan in a variety species, including in rodents, zebra finches and humans (Briga, 2016; Miller et al., 2002; Prospective Studies Collaboration, 2009). Its aging is widely observed (Douhard et al., 2017) and its decline is associated with behavioral and physical deterioration including foraging efficiency (Catry et al., 2006), loss of muscle mass (sarcopenia) and muscle strength (Colman et al., 2008; Sayer et al., 2008) and loss of body fat (Kuk et al., 2009). Hence body mass is a key trait associated with organismal functioning and survival in many species.

We here use an outdoor-living captive population of zebra finches exposed to natural weather fluctuations and monitored from birth till natural death (Briga et al., 2017; Briga and Verhulst, 2015a). These birds showed two levels of variation in lifespan. The first level is induced by an experimental manipulation of environmental quality. The environment can affect lifespan at all ages, but the development phase is thought of as a particularly important for adult lifespan and health (Lindström, 1999; Lummaa and Clutton-Brock, 2002; Metcalfe and Monaghan, 2001). In our study, we altered developmental conditions by cross fostering chicks to either small or large broods. Growing up in large broods impairs growth and hence, large broods represent a harsh environment (Briga et al., 2017; Griffith and Buchanan, 2010). However, the long-term effects of developmental conditions on lifespan can depend on the environmental conditions in adulthood (Bateson et al., 2004; Hanson and Gluckman, 2014). We thus experimentally manipulated foraging costs during adulthood and exposed birds from both developmental conditions to either low or high foraging costs in a 2×2 design. We further abbreviate the high foraging cost group as harsh (H) and the low foraging cost group as benign (B). Hence, we have four treatment combinations (BB, HB, BH, HH). The group that experienced a harsh environment during development and in adulthood (HH group) lived 6 months (12%) shorter compared to all other treatment combinations. Furthermore, females lived one month shorter than males (Briga et al., 2017). Thus, if aging trajectories are scaled to lifespan, we expect an accelerated onset and/or rate of body mass decline in females relative to males, and in the HH group relative to all other treatment combinations.

## Material & methods

### Experimental setup

The birds were reared in either experimentally small broods (with 2 or 3 chicks, modal =2) and large broods (between 5 and 8 chicks, modal=6). These brood sizes are within the range observed in wild (Zann, 1996). Growing up in large broods impairs growth (Briga, 2016; Briga et al., 2017). After nutritional independence and before the start of the foraging cost manipulation, i.e. between 35 days till approximately 120 days, young were housed in larger indoor cages with up to 40 other young of the same sex and two male and two female adults. Once adult, birds were subject to a long-term foraging experiment, (Koetsier and Verhulst, 2011). Briefly, birds were housed in eight single sex outdoor aviaries (LxHxW 310×210×150 cm) located in Groningen, the Netherlands (53° 13’ 0” N / 6° 33’ 0” E). Food (tropical seed mixture) water, grit and cuttlebone were provided *ad libitum*. In addition, the birds received fortified canary food (‘‘egg food”, by Bogena, Hedel, the Netherlands) in weighed portions. Each aviary contained an approximately equal number of birds and to keep densities within aviaries within a limited range, new birds were added regularly to replace those that died. The first batch was 3-24 months old when the experiment started and birds added later were 3 to 4 months old.

### Lifespan estimates

Group-specific estimates of median lifespan were taken from (Briga et al., 2017). In brief, there we estimated lifespan using two approaches, Cox proportional hazards (Cox, 1972) and Gompertz fits (Gompertz, 1825). Both approaches showed that (i) the median lifespan of the HH group was 6 months (12%) shorter relative to all other treatment combinations and (ii) females lived one month shorter than males. The environmental effect on lifespan was more pronounced in females than males (table 1) but this difference was not significant (Briga et al., 2017).

**Table 1:** Median lifespan and description of the changes with age and onset of body mass aging per experimental group and per sex. Median lifespan values are taken from (Briga et al., 2017)

| **Developmental environment** | **Benign** |  | **Harsh** |  |
| --- | --- | --- | --- | --- |
| **Adult environment** | **Benign** | **Harsh** | **Benign** | **Harsh** |
| **Males** |  |  |  |  |
| N | 75 | 75 | 73 | 81 |
| median lifespan [years] | 4.3 | 4.0 | 3.6 | 3.6 |
| aging trajectory | quadratic | quadratic | quadratic | quadratic |
| onset of decline [years] | 8.2 | 3.7 | 3.4 | 3.9 |
| **Females** |  |  |  |  |
| N | 68 | 75 | 73 | 77 |
| median lifespan [years] | 3.7 | 4.0 | 3.5 | 2.7 |
| aging trajectory | quadratic | quadratic | linear decline | linear increase |
| onset of decline [years] | 2.7 | 3.3 | 0.3 | no decline |

### Data collection

Between December 2007 and December 2015, we collected 15443 mass measurements on 597 individuals, with birds being measured between 1 and 95 times over their lifetime (Fig. S1A). Data were collected from individuals covering an age range from 0.4 months till 9.4 years (Fig. S1B) and at almost monthly intervals (Fig. S2B). Measurements were randomized across experimental groups. At the average age of 120 days (SD: 29 days), we measured body size, using the average of the tarsus and the headbill after transforming both to a standard normal distribution.

### Statistical analyses

Data for all traits were collected at any hour of the day and throughout the year (Fig. S2A & B). To avoid confounding age patterns with daily or seasonal effects, we corrected for daily and seasonal variation in trait values. To this end, we first investigated for each trait how best to correct for daily and seasonal variation in trait values (Supp. Information 2). Model selection approach (see below) indicated that we best captured daily and seasonal variation in 3 variables: (i) daylength, (ii) photoperiod dynamics (increase vs. decrease), (iii) time of measurement and their interactions (Table S1). Hence to obtain unbiased age estimates we use body mass values adjusted for daily and seasonal fluctuations in all further analyses.

Population level associations between trait values and age can be composed of two processes: (i) a within individual change in trait value with age and (ii) a between individual change due to selective mortality of individuals with certain trait values. We distinguished the contributions of these two processes using a within subjects centering approach (van de Pol and Verhulst, 2006; van de Pol and Wright, 2009). In this approach the within individual changes are captured in a Δage term, which is the age at measurement mean centered per individual. Within individual changes can also show terminal changes before death. We therefore added a terminal term as a separate variable, coded as a binomial factor for whether or not an individual died within the year following the measurement. The between individual change is captured by the term lifespan, mean centered across our population. For censored birds, i.e. those still alive (N=179) or that died an accidental death (N=16), lifespan is unknown and thus received a lifespan of zero. In this way, these birds contribute only to the estimate of within individual trait change while having no effect on the estimates of between individual trait change. To test whether within individual change is environment specific we included the interaction between Δage terms and our experimental manipulations. Tests for context dependent developmental effects were done with three-way interactions (e.g. Δage*development*adult).

All analyses were done using a general linear mixed modeling approach with the function ‘lmer’ of the package ‘lme4’ version 1.1-10 (Bates et al., 2015) in R version 3.2.1 (R Core Team, 2014). Experimental treatments during development (small vs. large broods), adulthood (low vs. high foraging costs) and their interaction were included as categorical variables. All analyses included individual as a random intercept and Δage and Δage^2^ nested within individuals as a random ‘slope’. The random slope quantifies the within-individual variation in aging and is required for the correct estimation of confidence intervals when investigating within individual changes (Schielzeth and Forstmeier, 2009). Such models require considerable sample sizes to accurately estimate fixed and random effects and our data (Fig. S1A & B) fulfilled those requirements (van de Pol, 2012). Residuals of all final models were normally distributed and without outliers (Figs. S4). Confidence intervals of model parameters were estimated with the Wald approximation in the function ‘confint’. In selected cases we report effect sizes, estimated as the ratio of the coefficient to the variable’s standard deviation, following equation 1 in (Nakagawa and Cuthill, 2007).

To find the model best supported by the data we used the model selection approach proposed by Burnham and Anderson (Burnham and Anderson, 2002; Burnham et al., 2011) based on second order Akaike Information Criterion (AICc) with the function ‘dredge’ of the package ‘MuMIn’ version 1.15.1 (Barton, 2009). In brief, this is a hypothesis-based approach that generates, given a global model, subset models that best fit the data. Better fitting models are indicated by their lower AICc and, as a rule of thumb, a decrease of 2 AICc is considered significant (Burnham and Anderson, 2002; Burnham et al., 2011).

## Results

We collected 15.443 measurements on 597 birds covering an age range from 0.4 months till 9.4 years (Figs. S1A & B). On average, birds reared in large broods weighed 0.56g (95%CI: −0.77, −0.35) less than birds reared in small broods (Table S2; ΔAICc=−22.4), and birds in the harsh adult environment weighed 0.66g (95%CI: −0.87, −0.45) less than birds in the benign adult environment (Table S2; ΔAICc=−32.4). The effects of both manipulations on mass were additive (Table S2; developmental * adult environment ΔAICc=+3.3; Fig 2). Because mass is to a large extent determined by an individual’s body size (r=0.56), we investigated to what extent the manipulation effects on mass were independent of size. Growing up in large broods resulted in smaller adult body size (N=594 individuals; t=−4.37; p=0.00001), while there was no association between the adult manipulation and size (t=−1.41; p=0.16). When we analyzed the manipulation effects on mass including body size as a covariate, we found that the effect of the brood size manipulation on mass approximately halved from 0.56 to 0.27g, but the effect of brood size on mass remained clear (ΔAICc=−1.7). In contrast, correcting for size made the adult manipulation effect on mass more evident (ΔAICc=−40.8; Fig. S3; see SI 3 for details). Thus, both manipulations affected mass, but the effect of the developmental manipulation was partially mediated via body size, while, as expected, the effect of the adult manipulation was size independent.

**Fig. 2:**
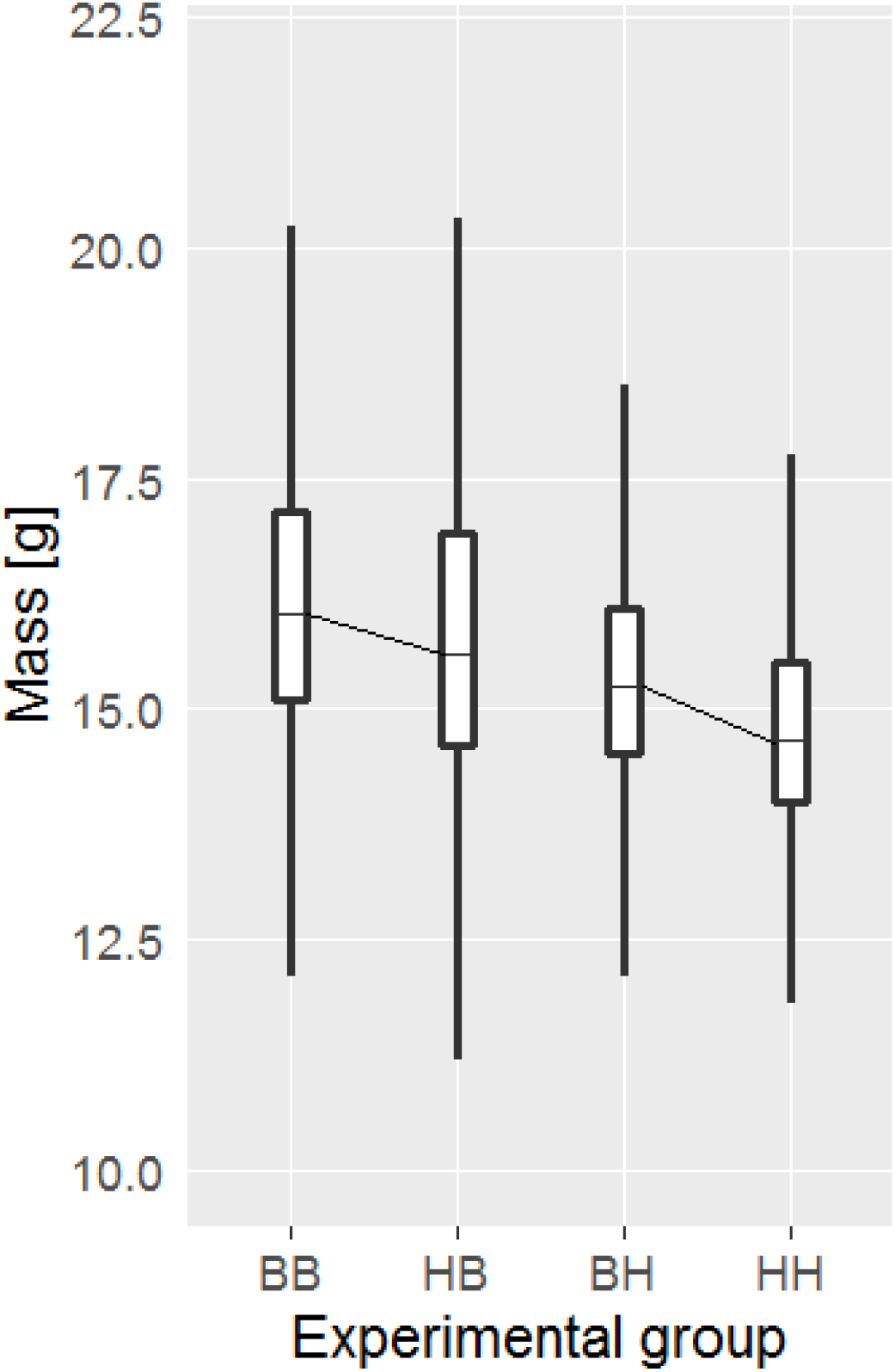
Harsh environments decreased mass. Shown are boxplots with median, quartiles and 95% CI. Statistical analysis showed the effects of developmental and adult environments to be additive for mass. Horizontal lines connect groups from different brood sizes in the same foraging treatment.

### Aging trajectories and lifespan between groups

We investigated the body mass aging trajectory within individuals in the different environmental treatment categories, testing for the scenarios in Fig. 1. In the complete dataset, the aging trajectory was best described by a quadratic shape, rather than a linear shape (ΔAICc=+375.5). A quadratic shape also fitted the data better than a terminal effect either on its own (ΔAICc=+354.5) or in combination with a quadratic or linear decline (+5.3<ΔAICc<+341.9). However, sexes differed in their age trajectory (Δage^2^ * sex ΔAICc=−46.2) and in their environmental susceptibility of the age trajectory (Δage^2^ * foraging treatment * sex ΔAICc=−37.9). Moreover, females were slightly heavier than males (not corrected for their size; ΔAICc=−50.7; Table S3). To gain better insight in these interactions, we further analyzed the sexes separately.

In males, the best fitting aging trajectory was quadratic (ΔAICc<−163.0; Table S4) independent of the environmental manipulations (ΔAICc>+7.1; Fig. 3C-F; Table S4). The quadratic random term varied little between individuals relative to individual as random intercept (variance explained: 1.8% vs. 81%), showing that individuals differed more in their mean mass than in their aging trajectory. A quadratic age term can reflect a trajectory that first increases and then decreases (or vice versa), but can also reflect e.g. a levelling off with increasing age. To discriminate between these patterns, we tested whether mass changed significantly with age pre- and post-peak. For all treatment combinations pooled, maximum body mass was reached at the age of 4.2 years. In the pre-peak phase, mass increased significantly with age (0.07 g/yr; 95%CI: 0.04, 0.11; ΔAICc=−8.0), and mass decreased post-peak, albeit not significantly (−0.03 g/yr; 95%CI: −0.17, 0.09; ΔAICc=+5.3). Thus, for all treatment combinations pooled, male body mass aging trajectories was quadratic, increasing till the age of 4.2 years followed by a non-significant decline.

**Fig. 3:**
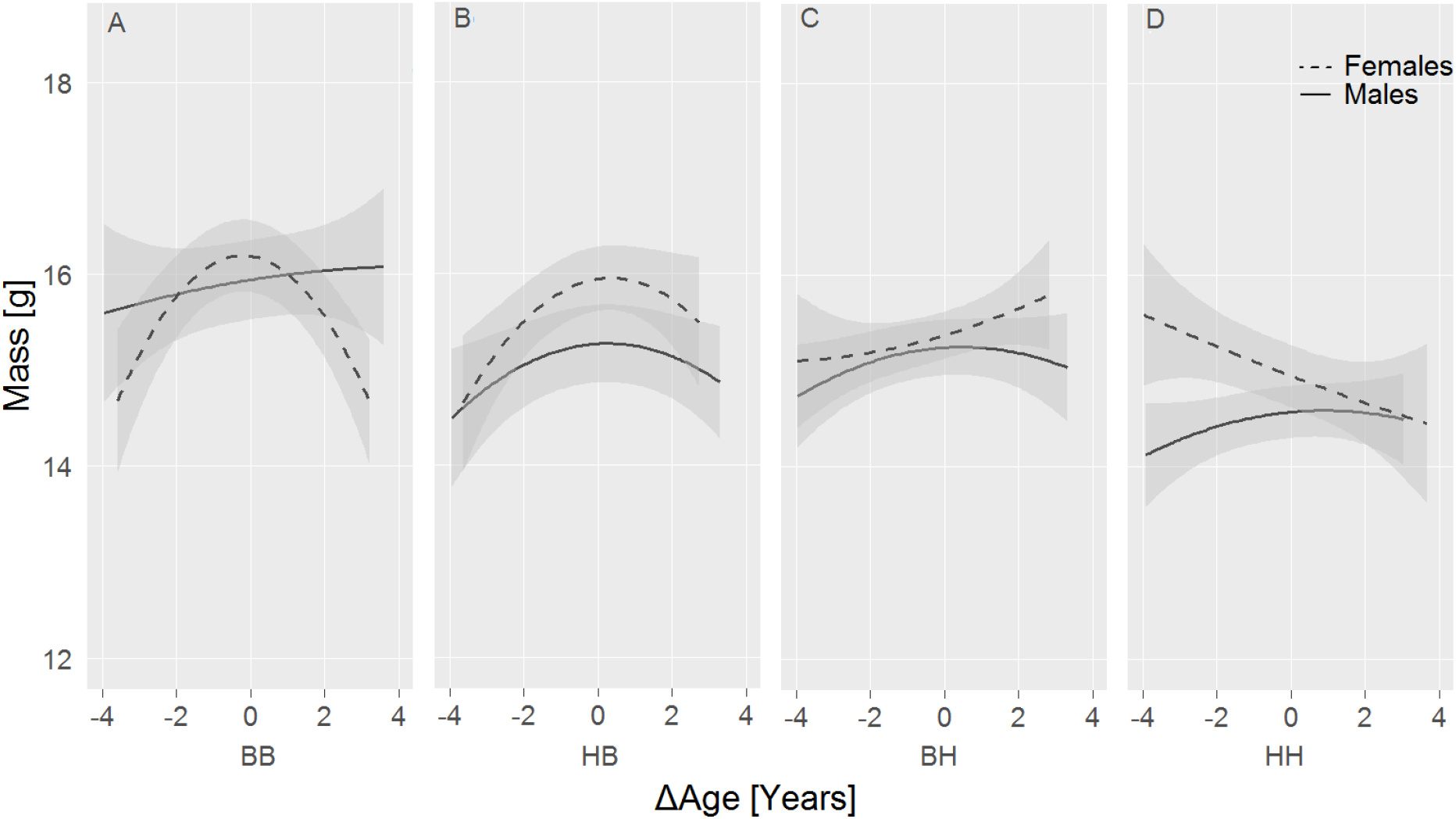
Within-individual mass age trajectories are sex and environment specific. Males show a quadratic age trajectory which shape was consistent across experimental groups (A-D). In contrast, (A & B) females showed a quadratic age trajectory when foraging costs were low but (C & D) when foraging costs were high the age trajectory was linearly which slope depended on developmental conditions: increasing and decreasing for birds from benign vs. harsh developmental conditions respectively. For data plots, see fig. S4.

To compare the scaling of the onset of body mass decline with the median lifespan of all treatment combinations, we analysed the aging trajectories of all treatment combinations in separate models. This more sensitive approach confirmed a quadratic aging trajectory for all treatment combinations (Fig. 3). However, for the BB group, male body mass peaked late at the age of 8.2 years (Fig. 3A). In the other treatment combinations body mass peaked earlier, at 3.7, 3.4 and 3.9 years for HB, BH and HH respectively (Fig. 3B-D). Given a median lifespan per group of 4.3, 4.0, 3.6 and 3.6 years for BB, HB, BH and HH respectively (Table 1), this shows that aging and lifespan are not scaled. Thus, analyzing all treatment combinations separately confirmed a quadratic aging trajectory for male body mass and the onset of the post-peak decline was not scaled to the group’s median lifespan.

In females, the aging trajectory differed between the benign and harsh foraging environment (Δage^2^ * treatment ΔAICc=−16.1; Table S5). Analyzing these treatments separately revealed that females in benign foraging environment had a quadratic trajectory (ΔAICc=−9.3; Fig. 3A & B), which was independent of brood size (ΔAICc>+6.5; Table S6A). The quadratic random ‘slope’ varied little between individuals relative to individual as random intercept (variance explained: 1.9% vs. 70%), showing that also female mass differed more between individuals in mean body mass than in the aging trajectory. Females reached their maximum mass at a younger age than males, at Δage = 0.03 years or an age of 3.2 years. The pre-peak increase in mass was four times larger than in males (0.29 g/yr; 95%CI: 0.18, 0.39; ΔAICc=−17.0). The post-peak decrease was between two and three times larger than the decrease in males, but not significant (−0.08 g/yr; 95%CI: −0.17, 0.01; ΔAICc=+3.5). Thus, for females in the easy treatment, mass changed quadratically with age, characterized by a steep increase with a peak halfway through their life that is followed by a shallow non-significant decline.

For females in the hard treatment the mass age trajectories were linear (ΔAICc=−8.1; Fig. 3C & D) and differed between birds from small and large broods (Δage * brood size ΔAICc=−10.9; Table S6B). For females reared in small broods, mass increased linearly with age, while mass decreased with age in females from large broods (Fig. 3C & D; Table S6B). Rates of mass change (in absolute value) was close to 0.1 g/yr for both groups (small broods: 0.10 g/yr; 95%CI: 0.03, 0.17; large broods: −0.14 g/yr; 95%CI: −0.35, −0.13). Thus, for females in the harsh adult environment mass changed linearly with age in a direction that depended on the developmental conditions.

### Aging trajectories and lifespan within groups

The approach in the analyses above implicitly estimates an average aging trajectory for all individuals from a given group. However, there could be an association between lifespan and the aging trajectory within experimental groups. We tested this using the interaction between individual lifespan and within individual age terms (Δage, Δage^2^ and terminal year) and comparing the fit of the new model relative to the fits of the models above. For males, adding any of interactions between Δage or Δage^2^ with lifespan to the best fitting model in table S4 resulted in a poorer model fit (ΔAICc>+4.8, Fig. 4A-D). The same result emerged for females in tables S6-S7 (ΔAICc>+6.7, Fig. 4E-H). Thus, body mass of individuals with different lifespans within experimental groups did not show different aging trajectories.

**Fig. 4:**
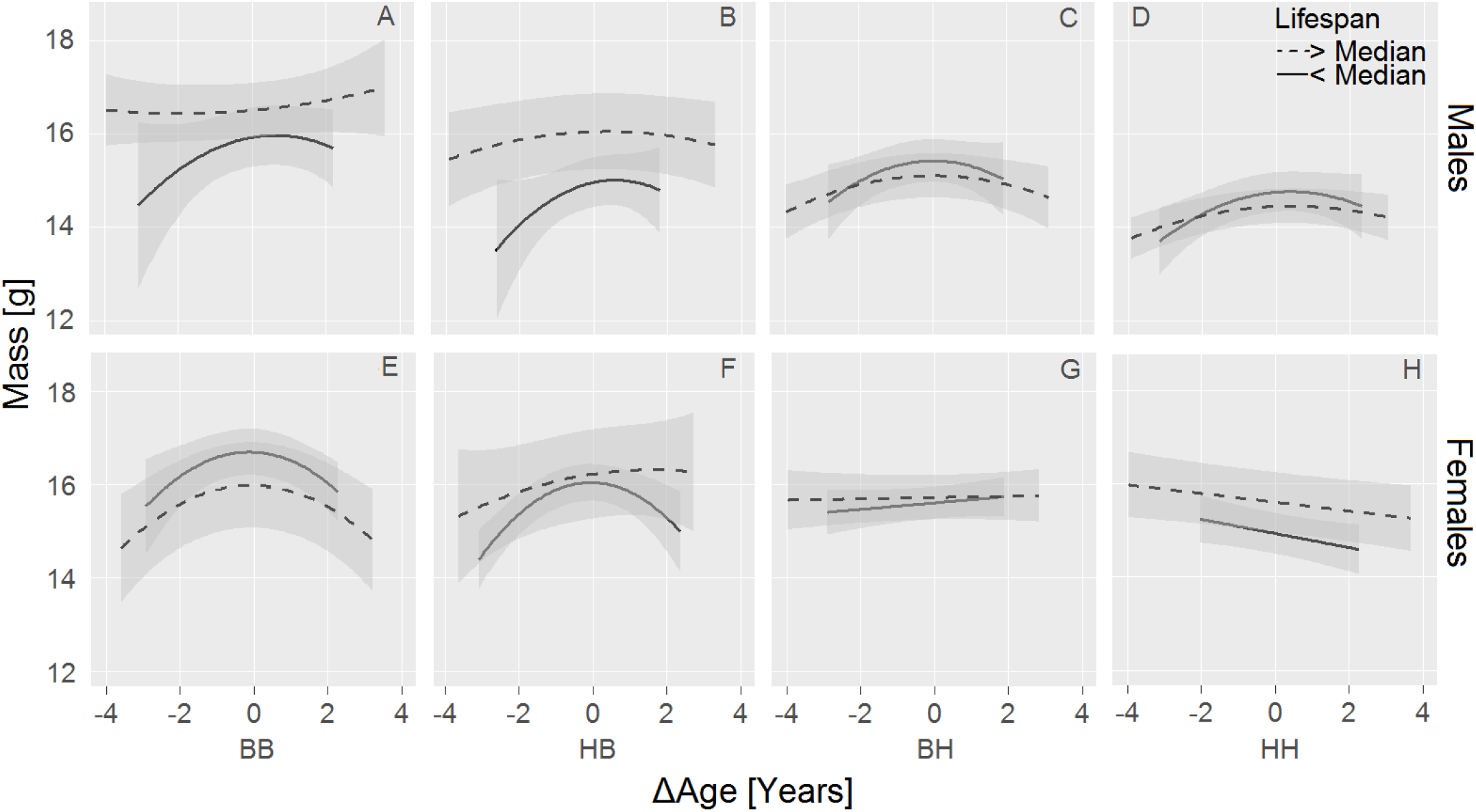
Within-individual aging trajectories for mass are independent of lifespan variation within experimental groups. For graphical purposes, age trajectories are shown for individuals living longer and shorter than median lifespan per experimental group. Analyses were done with lifespan as a continuous variable.

## Discussion

Identifying how phenotypes change with age (Fig. 1), what affects these changes and how they scale to lifespan are key questions in an ever longer living society. We here investigated whether environmental manipulations that shortened lifespan accelerated body mass declines in zebra finches. We found that male body mass increased with age, followed by a non-significant decline, and this was independent of experimental treatments during development and in adulthood. In females, the shape of the aging trajectory was treatment-specific: a quadratic shape as in males (but significant) in the benign adult environment (BB and HB groups), a linear decline for the BH group and a linear increase for the HH group. The environmental manipulations that shortened lifespan altered aging in females but not in males, but the effects on both traits were never scaled. This is partially because the environment can mold the shape of aging, a rarely studied phenomenon. However, females lived shorter than males and for most experimental groups, their body mass decline started earlier. Hence, environment-specific lifespan differences were not associated with body mass aging, but sex-specific differences were.

### Scaling of aging and lifespan

To what extent lifespan and aging are scaled remains poorly understood. A previous study using long-lived *C. elegans* mutants found that for several traits, aging started at the same age for long-lived mutants as for wild types (Bansal et al., 2015). Hence these long-lived mutants spent a larger proportion of their lives in an aged state. Whether such result can be extrapolated to more natural manipulations of lifespan remains an open question (Briga and Verhulst, 2015b). Our study shows that the longest-living groups spent proportionally more time decreasing body mass (table 1). Hence our results are consistent with the study of Bansal et al. 2015. It is unclear whether these results will apply to other traits. Within individuals, different traits age at different rates and shapes (Gaillard and Lemaitre, 2017; Hayward et al., 2015; Herndon et al., 2002), including in our zebra finches (Briga, 2016). This complexity can be seen in rapamycin experiments in rodents which extends lifespan and postpones aging of some traits, including body mass, but not of others (Fischer et al., 2015; Neff et al., 2013). Hence in our study, the longest-living groups spent a larger fraction of their lives with lower body mass, but to what extent this can be extrapolated to other traits and manipulations of lifespan requires further study.

### The shape of aging

For several experimental groups we found a quadratic body mass change with age. This shape is commonly observed in humans (reviewed in Kuk et al., 2009) and in laboratory rodents (Miller et al., 2002; Murtagh-Mark et al., 1995; Turturro et al., 1999; Yu et al., 1985). It was also observed in wild bighorn sheep *Ovis Canadensis* (Nussey et al., 2011) and in wild yellow-bellied marmots *Marmota flaviventer* (Kroeger et al., 2018). However, other aging shapes have also been reported: accelerating declines in Roe deer *Capreolus capreolus* (Nussey et al., 2011), terminal declines in Soay sheep *Ovis aries* (Hayward et al., 2015), accelerating and terminal declines in European badgers *Meles meles* (Beirne et al., 2015) and in male Alpine marmots *Marmota marmota* (Tafani et al., 2013). Note that most of these studies are in mammals. Previous data in captive zebra finches found that males gained weight with age, while females did not, but this was based on a small sample with three measurements per individual (Moe et al., 2009). Hence a variety of body mass aging shapes were described, largely biased towards mammals, of which a quadratic shape is the most abundant.

Population differences in body mass aging, possibly due to differences in environmental quality were found before, but these focused on the onset or rate of body mass declines (Douhard et al., 2017; Hämäläinen et al., 2014). Our study expands our view on the flexibility of body mass aging in previous studies by showing that not only the onset and the rate but also the shape of aging can be environment-specific. This has rarely been investigated as individual variation in the shape of aging trajectories is rarely tested for. Such variation can arise for example through canalization when the association between trait value and fitness is environment-specific (Boonekamp et al., 2018). The shape of aging is important though because it determines any comparisons in onset or rate of aging and any associations between aging and other traits such as lifespan. Thus, to what extent aging trajectories can differ between individuals for other traits and what determines this flexibility requires further study.

### Sex-specific aging

For those experimental groups with a quadratic shape, we found that aging started earlier in the shortest living sex, i.e. females. Theory on sex-specific aging predicts that the shortest living sex also ages fastest (Bonduriansky et al., 2008; Maklakov and Lummaa, 2013) and hence our result is consistent with the expectation. Sex-specific body mass aging was found in several wild mammals and several of these found results consistent with the expectation (Tafani et al., 2013) (Beirne et al., 2015), but see (Hämäläinen et al., 2014). However, there are many exceptions. Most notably, in humans women typically outlive males but their age associated body mass decline starts a decade earlier (reviewed in Kuk et al., 2009).

Body mass aging was more sensitive to environmental conditions in females than in males. Sex biased environmental sensitivity is well known in many species, although its causes in birds remain unclear (Jones et al., 2009). In zebra finches, some studies found that females were more sensitive to developmental conditions than males (e.g. de Kogel, 1997; Martins, 2004), although this is not a general finding (Griffith and Buchanan, 2010). In our study, lifespan showed a trend in that direction, albeit not significantly (Briga et al., 2017). Thus, the female biased environmental sensitivity of the body mass aging shape is consistent with some trends or results in this and other zebra finch studies.

### Manipulation effects on mass

Birds reared in large broods had lower mass in adulthood, in agreement with earlier studies (reviewed in Griffith and Buchanan, 2010), and this effect was almost entirely due to their smaller structural size. Increased foraging costs during adulthood also resulted in lower mass, independent of size or rearing brood size. Size independent mass variation typically reflects variation in energy reserves, and theory predicts energy reserves to increase with increasing starvation risk (reviewed in Brodin, 2007). Starvation risk is higher in the high foraging cost treatment because it increases vulnerability to factors that increase energy needs (e.g. temperature) or impair foraging (e.g. illness). However, increased energy reserves also incur energetic costs (Hambly et al., 2004; Kvist et al., 2001; Schmidt-Wellenburg et al., 2008). This reduces optimal energy reserves, an effect which will also be stronger in the high foraging costs treatment because birds spend more time flying (Koetsier and Verhulst, 2011). The lower mass of birds experiencing higher foraging costs suggests that the birds weighed the energetic costs of carrying extra mass more than the decrease in starvation risk. This result is consistent with the findings in experiments with captive birds and mammals in which foraging costs were increased without changing predictability, which consistently resulted in lower mass (reviewed in Wiersma and Verhulst, 2005).

The functional significance of body mass aging trajectories remains poorly understood. Possible body composition changes underlying the mass age trajectory include loss of body fat and skeletal muscle (Ballak et al., 2014; Kuk et al., 2009). While age related changes in body composition in birds are poorly known, body mass changes likely partially reflect changes in energy reserves. We here suggest two possible options for the aging trajectories. First, we found that there is an increase with age in mass-adjusted standard metabolic rate (SMR), i.e. the minimum energy expenditure of a post-absorptive adult animal measured during the rest phase at temperatures below thermoneutrality (Briga, 2016). Such a larger energy turnover also requires larger energy reserves. Second, changes in body mass indicate an age associated change in the balance between the benefits and costs of carrying the weight in energy reserves. The lower mass in older birds indicates that these birds weighed the energetic costs of carrying extra mass more at the cost of starvation risk. This will likely make them more vulnerable to abiotic and biotic and challenges including harsh weather conditions and disease.

